# Duplicating At-Risk Breadfruit (*Artocarpus* spp.) Accessions Using Air-Layer Propagation

**DOI:** 10.64898/2026.02.22.707215

**Authors:** Kaitu Erasito, Noel D. Dickinson, Tiffany M. Knight, Mike Opgenorth

## Abstract

Breadfruit (*Artocarpus altilis* (Parkinson) Fosberg) is a culturally and nutritionally significant perennial crop of the Pacific Islands. National Tropical Botanical Garden’s Kahanu Garden (Maui, Hawai‘i) maintains a breadfruit collection representing more than 150 traditional varieties, some unique or irreplaceable and requiring safety duplication to safeguard genetic diversity. However, aging trees exhibit variable vigor, potentially limiting clonal propagation outcomes. We assessed air layering as a strategy for conservation duplication, conducting 163 air-layer attempts across 26 priority accessions. We evaluated the influence of tree vigor, age, and branch characteristics on rooting success and survival to out-planting. Overall, 17% successfully rooted and 75% of those survived to out-planting, resulting in successful duplication of 16 of 26 at-risk accessions. Rooting success differed among vigor classes (33% for high-vigor trees; 11–16% for normal and feeble trees) and increased modestly with source tree age, while survival to out-planting declined with increasing age. Branch length and fruiting season were not associated with outcomes. These findings indicate that air layering can support conservation propagation in living collections, but success is strongly influenced by source tree age and condition. Initiating safety duplication while trees are physiologically robust is likely to improve long-term conservation outcomes.

## Introduction

Breadfruit (*Artocarpus* spp.) is a perennial tree crop central to Pacific Island food systems and biocultural heritage (Ragone 2007; Whistler 2009). Long histories of selection and exchange across island networks have generated exceptional morphology and genetic diversity, reflected in the many named varieties maintained in farmers’ plantings and living collections (Kaoh *et al*. 2026; Zerega *et al*. 2015). While breadfruit origins and evolutionary pathways continue to be refined, its importance to food security, cultural continuity, and diversified agroecosystems across Oceania is well established (Berning *et al*. 2022; PIFON 2020; Ragone 2007).

Conservation of breadfruit diversity relies heavily on vegetative approaches because many varieties are seedless and seeded types may not produce true-to-type progeny (Ragone 2007). For this reason, living collections play a critical role in safeguarding breadfruit genetic resources, particularly for culturally significant and rare crop varieties (Dempewolf *et al*. 2023; FAO 2019; Khoury *et al*. 2022). The Breadfruit Institute at the National Tropical Botanical Garden (NTBG) curates one of the world’s most comprehensive breadfruit conservation collections, including more than 200 accessions at Kahanu Garden, Maui. Many trees were planted decades ago and now exhibit varying vigor and increased vulnerability to stressors, heightening the risk of loss of unique accessions (Fig. 1A–B).

**Figure 1.**
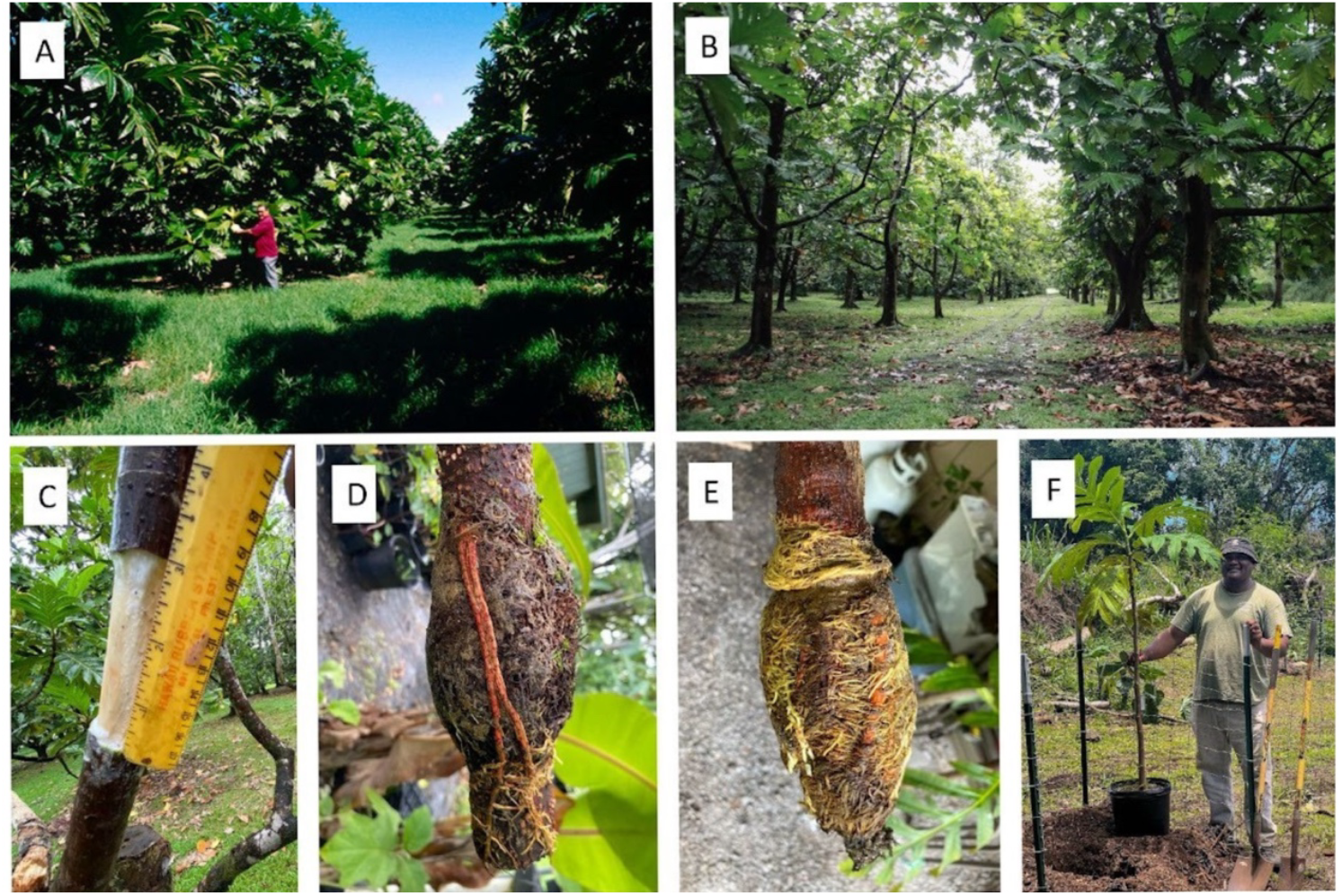
Air-layer propagation of at-risk breadfruit (*Artocarpus* spp.) at Kahanu Garden National Tropical Botanical Garden (NTBG), Maui, Hawai‘i. **(A–B)** Breadfruit living collection and representative source trees within the collection. **(C)** Measurement of branch diameter and girdling incision with removal of a section of bark. **(D–E)** Successfully rooted air layers showing varying levels of root development and coverage. **(F)** Potted air-layered tree prepared for out-planting. The individuals depicted in this figure are the authors of this study, who consent for publication. Photographs: **(A–B)** Diane Ragone, **(C–E)** Kaitu Erasito, **(F)** Mike Opgenorth.

Reliable clonal propagation methods are essential to maintain varietal identity, genetic integrity, and preventing irreversible loss (Ragone 2007; Whistler 2009). Traditional approaches in the Pacific include harvesting root shoots (suckers) and root cuttings, but in dense repository plantings, intertwined root systems can complicate confirmation of varietal identity. Other methods (e.g., grafting, tissue culture) may require specialized infrastructure or carry risks of physiological or genetic variation (Murch *et al*. 2008; Solomon *et al*. 2021).

Air layering (marcotting) is a low-technology field method that induces roots on intact branches prior to detachment, enabling clonal propagation without root excavation (Bridgemohan *et al*. 2016; Leakey 2014). However, source tree age may influence vegetative propagation outcomes through multiple physiological pathways, including carbohydrate availability, hormonal balance, and cambial activity (Davies *et al*. 2018; Leakey 2014). In many woody species, adventitious rooting capacity declines with ontogenetic aging (Hartmann *et al*. 2011). Because air layering relies on localized adventitious root formation, physiological variation occurring along the branch axis may also influence success. Branch segments positioned closer to metabolically active tissue (apical meristem and foliage) may provide more favorable conditions for vegetative propagation outcomes (Hartmann *et al*. 2011; Leakey 2018). Additionally, propagules derived from older trees may exhibit altered post-detachment resilience or establishment capacity due to changes in physiological condition (García *et al*. 2023; Qui *et al*. 2021). Although these factors are well studied in other fruit tree species, to our knowledge, the specific roles of tree vigor, age, and branch length on air layering success have not been examined for breadfruit and in living collections contexts.

This study assessed air layering as a tool for safety duplication of aging and at-risk breadfruit accessions at Kahanu Garden (NTBG). We examined (1) whether source tree age was associated with vigor class, (2) whether tree age, tree vigor class and branch characteristics influence rooting success, and (3) whether source tree age and vigor influenced survival of successfully rooted air layers to out-planting. By identifying factors associated with successful and unsuccessful air-layer propagation across stages of establishment, this study aims to inform practical monitoring benchmarks and management strategies for safeguarding breadfruit genetic resources in living collections.

## Methods

### Study site and source trees

The study was conducted from 2022–2023 within the breadfruit collection at NTBG’s Kahanu Garden, in Hāna, Maui, Hawai‘i. The site is warm and humid year-round, with a distinct wet season from November to April (HCDP n.d.). We identified 26 priority accessions for safety duplication, including accessions representing a sole living tree of a named variety within the collection. According to NTBG living collections records, the selected trees ranged from 21 to 47 years in age at the time of air layer set (Appendix A Table 1) and included *A. altilis* (n = 23) and *A. altilis* ×*mariannensis* (n = 3). Tree vigor was assessed visually and classified as feeble, normal, or high. A detailed source tree list is provided in Appendix A.

### Air-layer propagation

Non-fruiting branches with healthy growth were selected. Branch length (cm; branch union to apical meristem) was measured prior to installation (Fig. 2A). A bark section (5–8 cm) was removed on semi-lignified branch tissue, typically 12–18 cm from the branch base (Fig. 1C; Fig. 2B). Rooting hormone was applied to the upper incision edge in most cases, the wound was wrapped with moist sphagnum moss, enclosed in plastic film, and covered with foil to exclude light. Air layers were monitored and harvested when root development stabilized (Fig. 1D–E). Air layers were set opportunistically primarily during peak fruiting months in Hawai‘i (July–November; Liu *et al*. 2014), limiting statistical evaluation of seasonal effects. Attempts showing rot, desiccation, or breakage were recorded as unsuccessful. A stepwise protocol is provided in Appendix B Figures 1–5.

**Figure 2.**
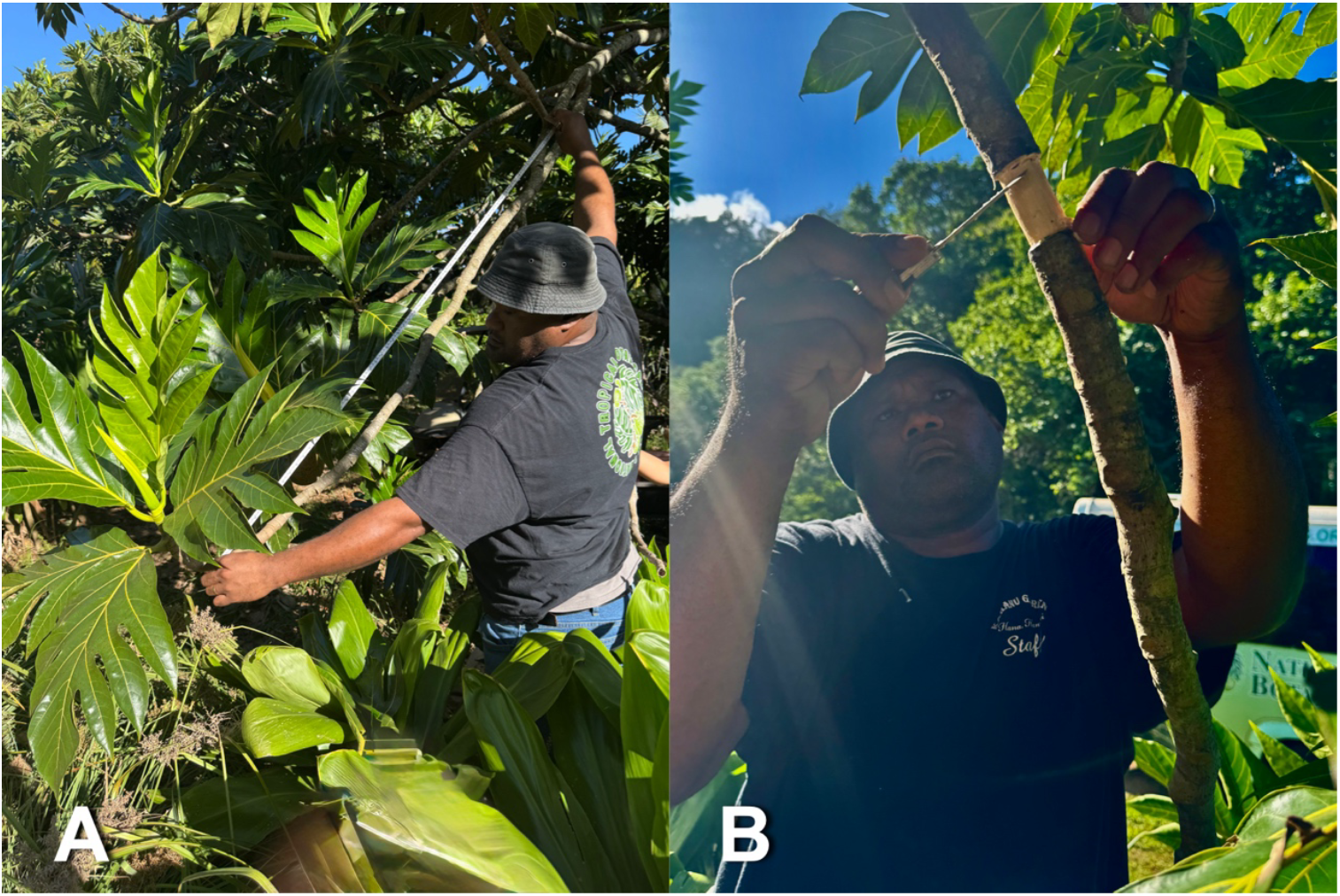
Selected branch measurement and girdling during air-layer propagation of breadfruit (*Artocarpus* spp.). **(A)** Measurement of selected branch from apex to base (branch union). **(B)** Removal of 2–3 cm band of bark and cambium just below a node to expose xylem, the area is scraped of cambial tissue prior to media application. The individuals in this figure are the authors of this study, who consent for publication. Photographs: Noel Dickinson.

### Statistical analysis

Analyses were conducted using R/RStudio (Posit Team 2023; R Core Team 2023), with graphics produced using ggplot2 (Wickham 2016). Means (± SD) are reported where appropriate. Associations between source tree age and vigor class (feeble, normal, high) were evaluated at the tree level (n = 26) using a Kruskal–Wallis’ rank-sum test. Rooting success (rooted vs. not rooted) was evaluated using binomial generalized linear models (GLMs), with source tree age and vigor class as predictors. Branch length was evaluated as an additional predictor of rooting success using binomial GLMs. Survival to out-planting among successfully rooted air layers was evaluated using binomial GLMs, with source tree age and vigor class as predictors; model terms were assessed using likelihood-ratio χ^2^ tests. Among successfully rooted air layers, relationships between days to root development and percent root coverage were evaluated using linear regression. Statistical significance was assessed at α = 0.05.

## Results

### Source trees

Source tree age among the 26 accessions ranged from 21 to 47 years, and trees were categorized as high-vigor (23%), normal-vigor (35%), or feeble (42%; Table 1). Tree age differed among vigor classes at the tree level (χ^2^ = 6.58, *P* = 0.037), with high-vigor trees older on average (mean 39.2 years) than feeble (33.6 years) or normal-vigor trees (30.4 years). Most air-layer attempts were conducted during peak fruiting months (July– November; n = 155).

**Table 1.** Summary of air-layer attempts, rooting success, and survival to out-planting by source tree vigor class for 26 priority breadfruit (*Artocarpus* spp.) accessions. Tree vigor was assessed visually and categorized as feeble, normal, or high. Values show total numbers of attempts, successfully rooted air layers, and air layers surviving to out-planting.

| Vigor class | Attempts | Rooted | Rooted (%) | Out-planted | Out-planted of rooted (%) |
| --- | --- | --- | --- | --- | --- |
| Feeble | 73 | 12 | 16 | 10 | 83 |
| Normal | 66 | 7 | 11 | 7 | 100 |
| High | 24 | 8 | 33 | 7 | 88 |
| **Total** | **163** | **32** | **17** | **24** | **75** |

### Air layer rooting success

Across 163 air-layer attempts, 32 successfully developed roots (17%; Fig. 3). Rooting success increased with tree age (binomial GLM, z = 2.55, *P* = 0.011; odds ratio ≈ 1.14 yr^-1^). Rooting rates differed among vigor classes in adjusted analyses (χ^2^ = 6.6, *P* = 0.037): 16% of attempts on feeble trees successfully rooted, 11% on normal-vigor, and 33% on high-vigor trees rooted (Table 1; Fig. 4). Yet vigor class was not independently associated with rooting success after accounting for age in a multivariable model (*P* = 0.77).

**Figure 3.**
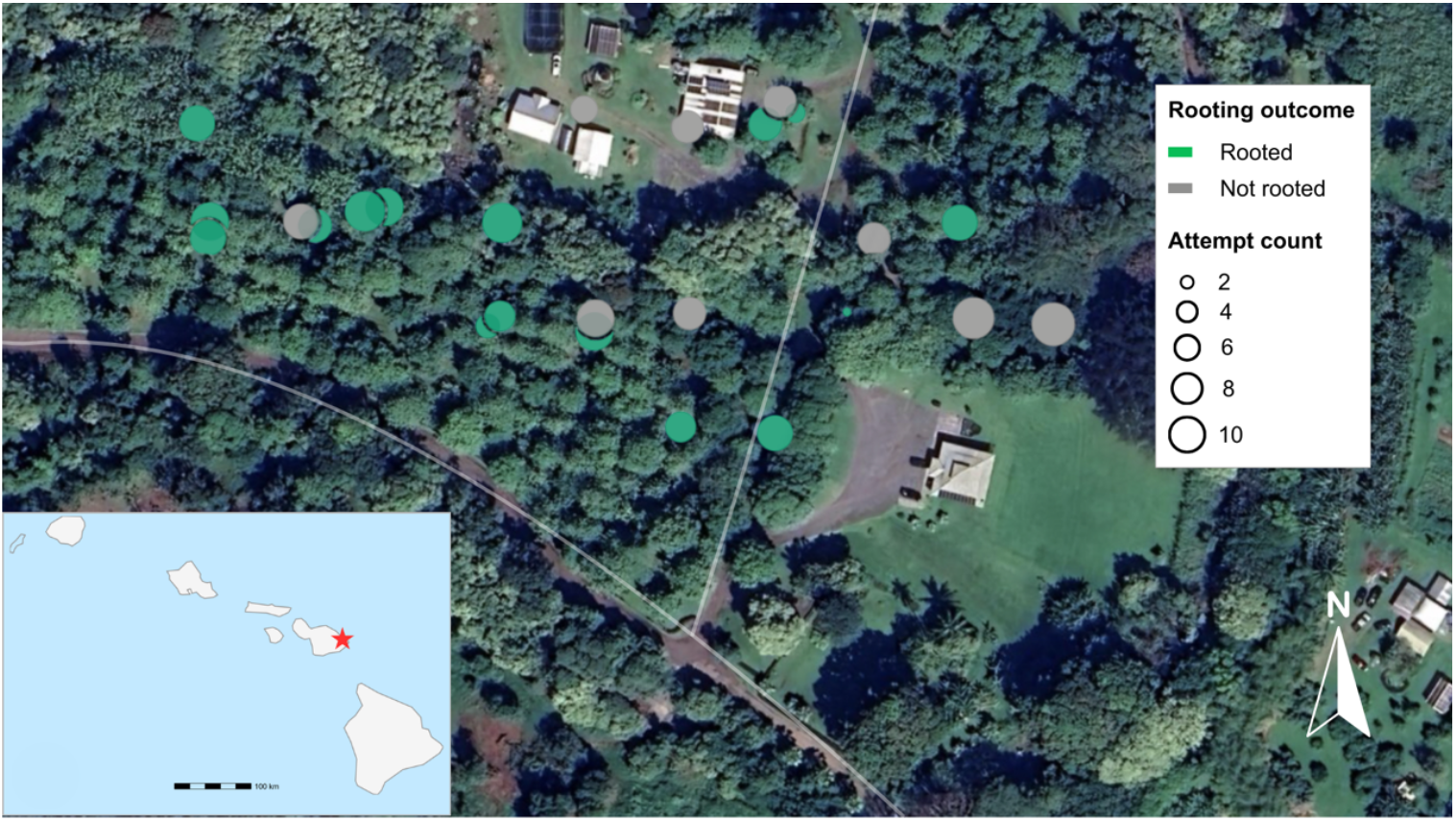
Map showing the location of breadfruit source trees included in the air-layer propagation study at Kahanu Garden (National Tropical Botanical Garden), Hāna, Maui, Hawai‘i. Each point represents a source tree, scaled by the number of air-layer attempts conducted per tree. Color indicates rooting outcome, with green symbols denoting trees from which at least one air layer successfully rooted. Inset map shows the location of Kahanu Garden within the Hawaiian Islands. Google Earth satellite image (Airbus 2023).

**Figure 4.**
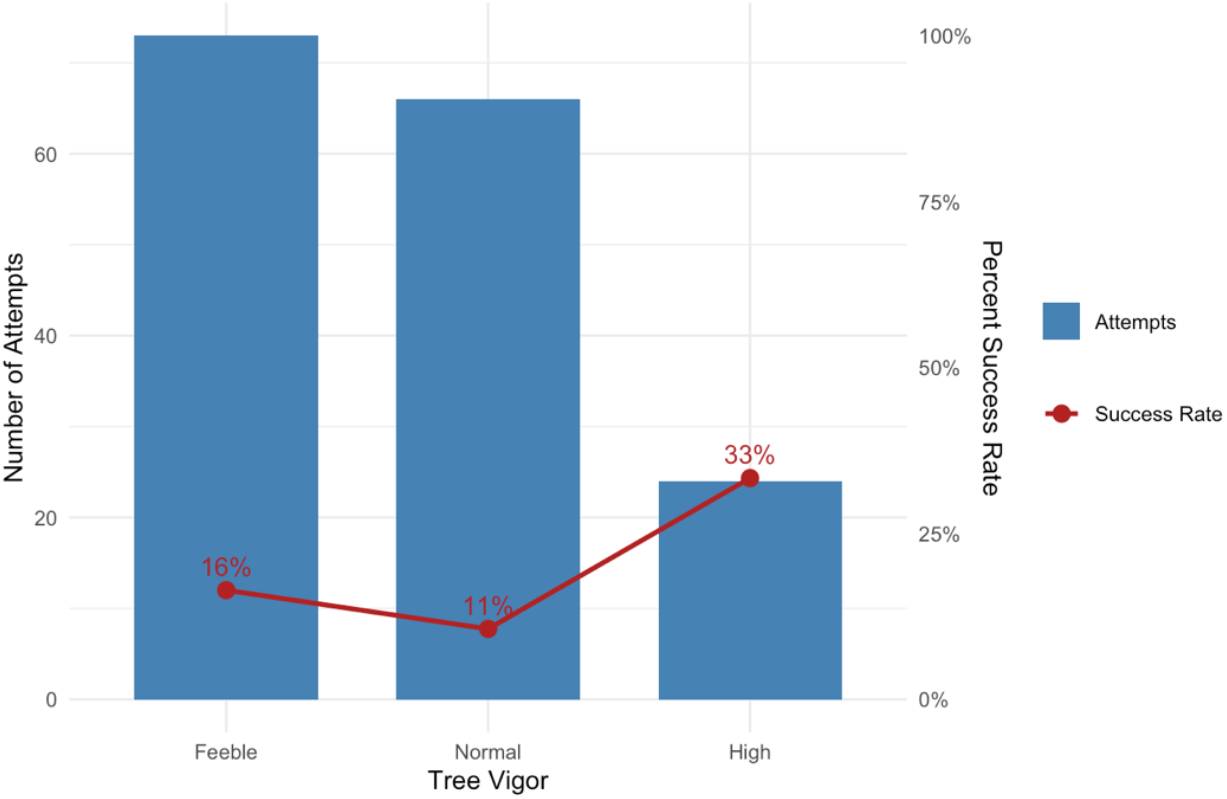
Number of air-layer attempts and root development rates by source tree vigor class (feeble, normal, high). Bars indicate the total number of air-layer attempts per vigor class, and points connected by a line indicate the proportion of air layers with successful root development, which differed among vigor classes.

Branch length among all attempts ranged from 36 to 305 cm (mean ± SD = 119 ± 44.8; median = 122) and overlapped substantially between successful and failed air layers (*P* = 0.39; Fig. 5). All successful air layers occurred within the bark removal range used (5–8 cm).

**Figure 5.**
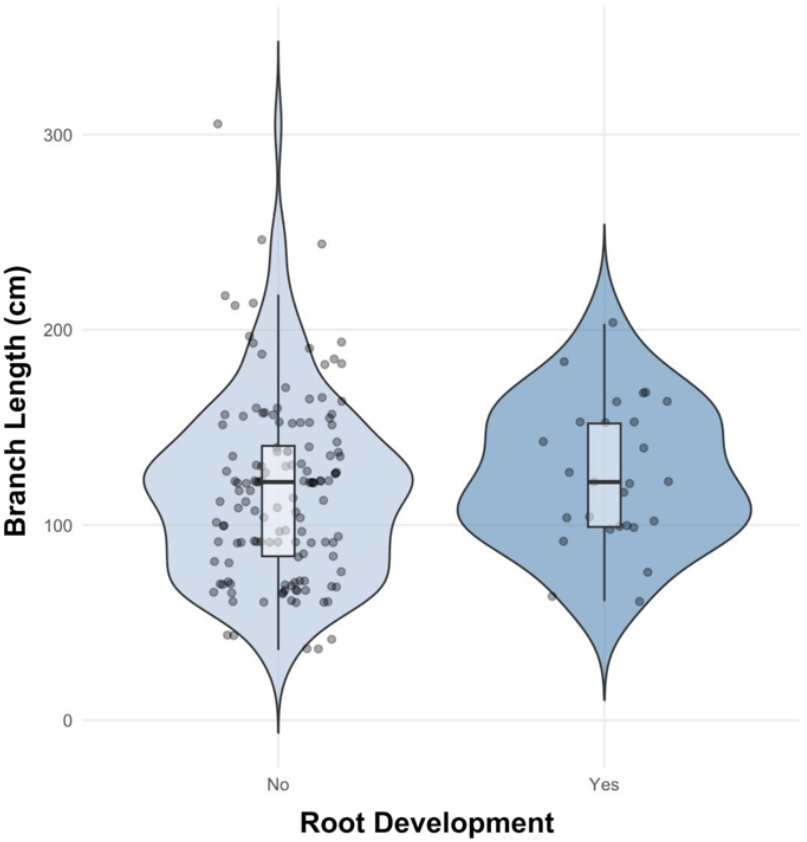
Distribution of branch lengths for successful and unsuccessful air-layer attempts. Violin plots show the distribution of branch lengths (cm) for air layers that failed to develop roots (“No”) and those that successfully rooted (“Yes”); points represent individual air-layer attempts and boxplots indicate median and interquartile range. Branch length did not differ significantly between rooted and non-rooted air layers.

Time to root development ranged from 55 to 212 days (mean ± SD = 122.3 ± 45; median = 114 days), with root development generally observed after 10–14 weeks. Among successfully rooted air layers (n = 32), percent root coverage declined modestly with increasing days to harvest (*R*^*2*^ *=* 0.17, *P* = 0.019; Fig. 6).

**Figure 6.**
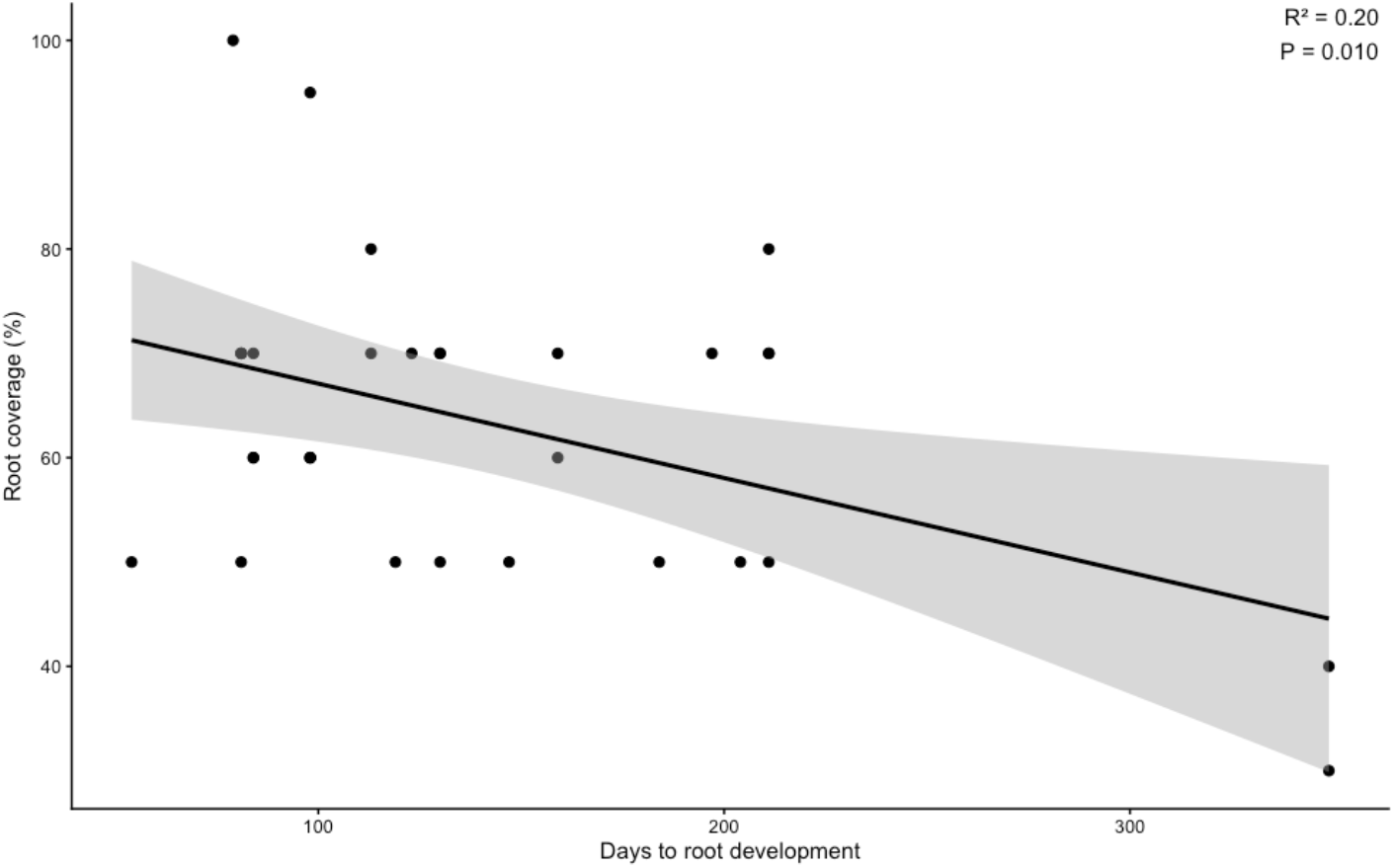
Relationship between days to root development and root coverage (%) among successfully rooted air layers (n = 32). Points represent individual air layers; the solid line indicates the fitted linear regression with 95% confidence interval (shaded). Root coverage declined with increasing time to root development (*R*^2^ = 0.20, *P* = 0.010).

### Successful air layer out-planting

Of the 26 at-risk breadfruit accessions, 16 were successfully duplicated and out-planted (Appendix A). Of rooted air layers (n = 32), 24 survived to out-planting (75%; Fig. 1F). In adjusted analyses, survival to out-planting declined with increasing tree age (z = –2.38, *P* =0.017), corresponding to an approximate 20–25% decrease in odds per additional year (odds ratio ≈ 0.76–0.80 yr^-1^). In multivariable models including vigor class, the negative association with age remained but was marginal (z = –1.86, *P* = 0.063), and vigor class was not independently associated with survival (*P* = 0.68).

## Discussion

This study evaluated air layering as a practical method for safety duplication of aging and at-risk breadfruit accessions in a living collection. Specifically, we found that (1) source tree age differed among vigor classes at the tree level, with high-vigor trees older on average; (2) rooting success increased modestly with tree age, whereas branch length was not associated with rooting success and vigor was not independently predictive after accounting for age; and (3) survival to out-planting declined with increasing age, independent of vigor.

### Vigor and age influenced different stages of success

Source tree vigor was associated with rooting performance, with much higher rooting rates observed among high-vigor trees. This is consistent with propagation literature linking physiological status to adventitious rooting capacity (Leakey 2014; Singh *et al*. 2004). Vigorous trees likely possess greater carbohydrate reserves, more favorable hormonal balance, and stronger hydraulic function to support root initiation and early growth following girdling. However, when tree age was included in the multivariable models, vigor was no longer independently predictive of rooting success. This suggests that chronological age accounted for much of the variation initially attributed to visible vigor class. Feeble trees, which are often those most urgently targeted for conservation duplication, exhibited lower unadjusted rooting rates in this study and may experience physiological constraints that reduce propagation responsiveness, creating challenges for repository management (Khoury *et al*. 2022; Tay *et al*. 2008).

Tree age influenced propagation outcomes at specific stages. Rooting success increased modestly with tree age, suggesting that older trees, in this collection, retained the capacity to initiate adventitious roots. Against expectations, high-vigor trees were older, and age differed among vigor classes, demonstrating that chronological age and visible condition were not inversely aligned in this context. However, among successfully rooted air layers, survival to out-planting declined with increasing source tree age. Although this association weakened slightly in multivariable models, the direction and magnitude were consistent, suggesting reduced establishment resilience for air layer propagules of older source trees. This contrasts with previous assessment of breadfruit in community-managed systems, where age was associated with vigor and structural characteristics (Dickinson *et al*. 2025), suggesting that age-related physiological change may not manifest uniformly across contexts.

The stage-specific pattern observed here — increased rooting but reduced post-detachment survival — may therefore reflect dynamics between initial regenerative responsiveness and longer establishment resilience. Such divergence aligns with broader horticultural observations of vegetative propagation performance variance across woody perennial lifespans (Leakey 2014; Singh *et al*. 2004). For living collections, these findings underscore the importance of initiating safety duplication before accessions enter advanced age or decline in vigor.

### Branch length was not limiting

Branch length did not predict success across the observed range. Provided that branches met minimum physiological and structural requirements, these findings suggest practitioners did not need to restrict air layer placement to a specific preferred branch length for higher success. This flexibility is important when working within constrained canopies or with aging trees exhibiting structural damage.

### Root development variability and practical benchmarks

Root development was typically observed within 10–14 weeks, consistent with prior breadfruit observations (Bridgemohan *et al*. 2016). Among rooted layers, longer time to harvest was modestly associated with lower percent root coverage, suggesting that extended intervals did not necessarily translate into greater root mass. This pattern likely reflects unmeasured drivers or microenvironmental factors influencing both root initiation and growth (Davies *et al*. 2018; García *et al*. 2023; Leakey 2014). Future work integrating physiological measurements could improve predictive accuracy and monitoring efficiency.

### Implications for living-collection management

Importantly, these findings reflect conservation duplication objectives rather than commercial multiplication. In conservation settings, air layering offers a low-technology option that can be implemented directly on source trees without root excavation or reliance on specialized infrastructure. Even modest success rates — duplicating 16 of 26 at-risk trees — may justify the effort when safeguarding culturally and genetically irreplaceable accessions, particularly where alternative collection or propagation pathways are not possible (Dempewolf *et al*. 2023; Khoury *et al*. 2022).

Our results provide empirical benchmarks for breadfruit air layering in a living collection and reinforce a clear management message: branch placement is less critical than overall source tree age and condition. Although older trees may initiate rooting successfully, propagules derived from older source trees exhibited reduced survival to out-planting. Safety duplication efforts should prioritize physiologically robust trees and avoid delaying action until decline is evident.

## Conclusion

Air layering can support safety duplication of breadfruit accessions in living collections, but success varies across propagation stages and is influenced primarily by source tree age and condition. In this study, 17% of air layers rooted and 75% of rooted layers survived to out-planting. Rooting increased modestly with source tree age, and although high-vigor trees exhibited higher adjusted rooting rates, vigor was not independently predictive once age was accounted for. In contrast, survival to out-planting declined with increasing source tree age. Branch length and seasonal timing were not associated with outcomes. These results provide practical benchmarks for conservation propagation and underscore the value of proactive duplication strategies to safeguard irreplaceable breadfruit genetic resources.

## Acknowledgements

This work was supported by the Ceres Trust, Omidyar Foundation, and Robert and Lien Chen Family Foundation, and by donors to the National Tropical Botanical Garden and the Breadfruit Institute. NTBG provided access to Kahanu Garden living collection, collections data, and staff support. We thank Kevin Houck for collection records assistance and the Kahanu Garden staff for field support. We are grateful to Diane Ragone for guidance on priority accession selection and to Nina Rønsted for early administrative leadership enabling project initiation. We also acknowledge Mike De Motta, Noa Lincoln, Dana Shapiro, and Mahele Farm for collaboration and support, and thank Dan Rudoy, Mariella Peck, Elliot Gardner, and Kelsey Rogers for contributions to project development, field support, and data collection.

