## Supplemental A Table 1 for "Duplicating At-Risk Breadfruit (*Artocarpus* spp.) Accessions Using Air-Layer Propagation"

**Appendix A. Table 1.** Priority accessions air layered. Columns: NTBG accession number, variety, taxon (Aa = *Artocarpus altilis*; Aam = *A. altilis* × *mariannensis*), island, tree age (years), vigor (+ = low; ++ = normal; +++ = high), attempts, rooted, and out-planted count.

| NTBG Accession | Variety | Taxon | Island Group | Tree Age | Vigor | Air Layers | | |
|---|---|---|---|---|---|---|---|---|
| | | | | | | Attempts | Rooted | Out-planted |
| 020498.001 | Blo lili | Aa | Solomon Islands | 21 | ++ | 7 | 0 | 0 |
| 020500.001 | Blo toki | Aa | Solomon Islands | 21 | ++ | 14 | 0 | 0 |
| 890154.001 | Hamoa | Aa | Society Islands | 35 | +++ | 1 | 0 | 0 |
| 900248.001 | Huero ninamu | Aa | Society Islands | 35 | ++ | 6 | 3 | 3 |
| 890162.002 | Lemai | Aa | Mariana Islands | 36 | + | 8 | 0 | 0 |
| 890184.001 | Luthar | Aam | Yap | 36 | + | 6 | 0 | 0 |
| 900259.001 | Malphang | Aa | Vanuatu | 35 | ++ | 8 | 1 | 1 |
| 900262.001 | Manua | Aa | Samoa | 35 | + | 9 | 1 | 1 |
| 900263.001 | Masee | Aa | Samoa | 35 | ++ | 8 | 1 | 1 |
| 890461.001 | Mei kakano | Aa | Marquesas Islands | 36 | +++ | 5 | 4 | 3 |
| 890466.002 | Mei koeng | Aam | Chuuk | 34 | + | 6 | 0 | 0 |
| 890463.002 | Patara | Aa | Society Islands | 35 | + | 7 | 0 | 0 |
| 900233.002 | Pulupulu | Aa | Rotuma | 35 | + | 7 | 1 | 1 |
| 770520.001 | Puou | Aa | Samoa | 47 | +++ | 2 | 2 | 1 |
| 890156.001 | Puou (Tahitian) | Aa | Cook Islands | 36 | + | 7 | 2 | 2 |
| 900234.001 | Samoan 1 | Aa | Fiji | 35 | + | 3 | 2 | 2 |
| 900261.001 | Samoan 2 | Aa | Fiji | 35 | + | 7 | 3 | 2 |
| 900281.001 | Tehelewa | Aa | Solomon Islands | 36 | +++ | 4 | 0 | 0 |
| 900281.002 | Tehelewa | Aa | Solomon Islands | 35 | ++ | 7 | 1 | 1 |
| 890456.001 | Toro | Aa | Solomon Islands | 36 | ++ | 6 | 0 | 0 |
| 890453.001 | Ulu afa | Aam | Tokelau | 36 | + | 8 | 1 | 1 |
| 770521.001 | Ulu ea | Aa | Samoa | 47 | +++ | 6 | 4 | 1 |
| 890155.002 | Ulu sina | Aa | Samoa | 36 | ++ | 9 | 1 | 1 |
| 770524.001 | Ulu tala | Aa | Samoa | 47 | +++ | 6 | 0 | 0 |
| 900226.001 | Undet CV 07 | Aa | Fiji | 36 | + | 5 | 4 | 2 |
| 890471.001 | Uto dina | Aa | Fiji | 36 | ++ | 1 | 1 | 1 |
| **Total** | | | | | | 163 | 32 | 24 |
