## Supplemental Data 1 for "Duplicating At-Risk Breadfruit (*Artocarpus* spp.) Accessions Using Air-Layer Propagation"

### Appendix B

The individuals depicted in this figure are the authors of this study, who provided consent for plublication.

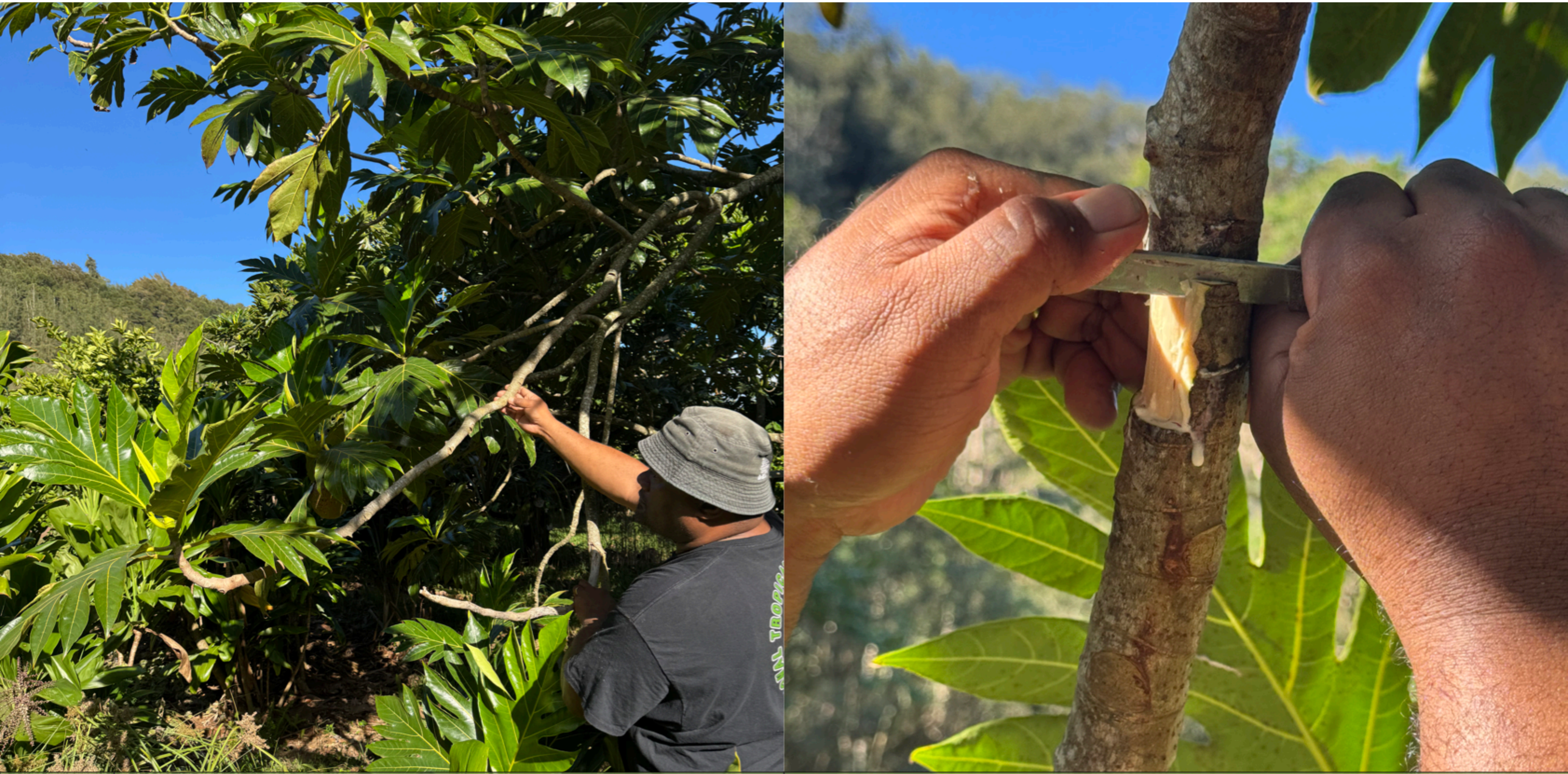

| Step | Procedure | Key considerations |
|---|---|---|
| 1 | Select a healthy, semi-hardwood branch. Identify node. Girdle branch just below node. | **Tools**: clean, sharp knife. **Avoid**: diseased, damaged, or branches with fruit load; damaging or removing node which may delay or prevent root initiation. |

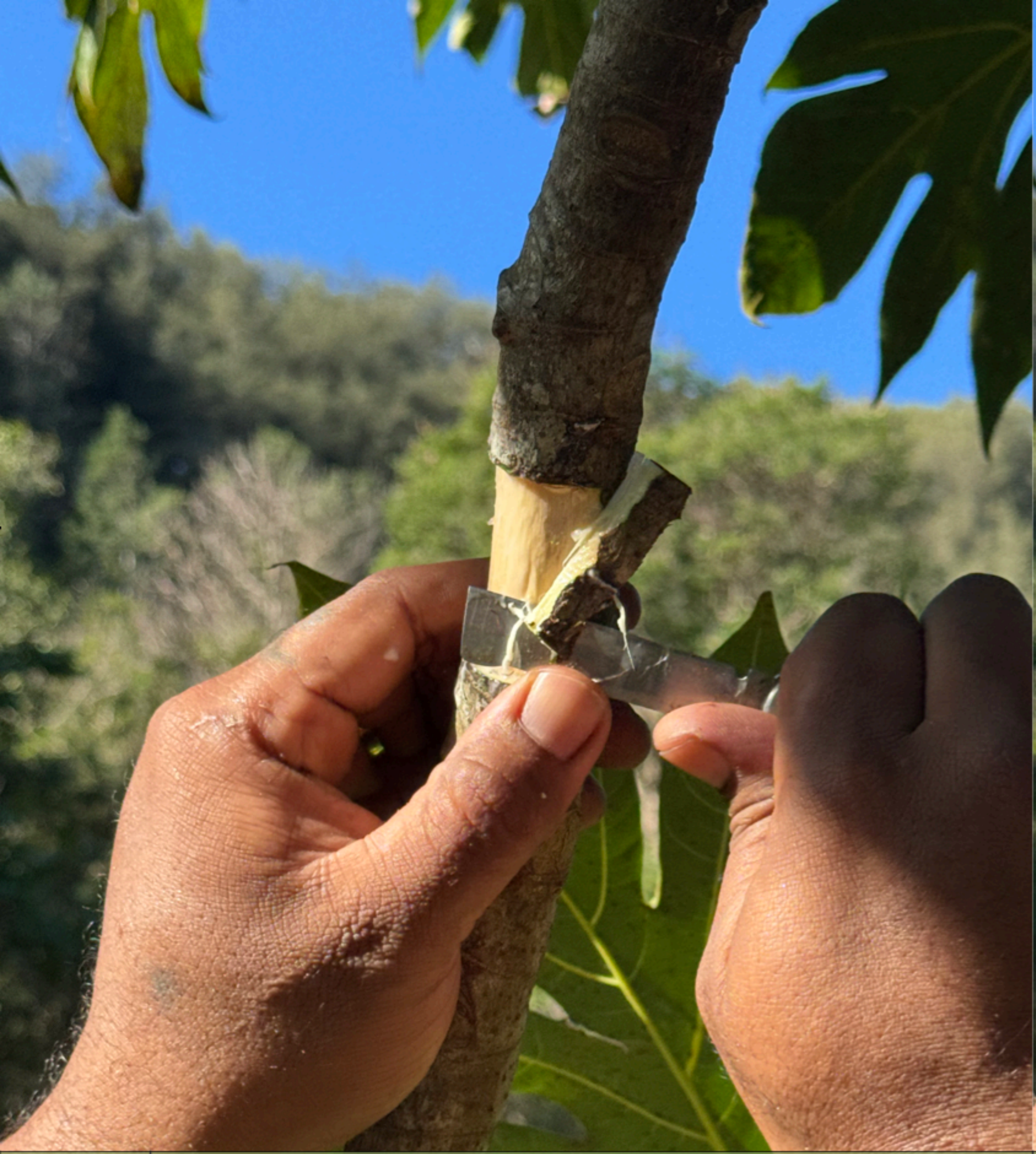
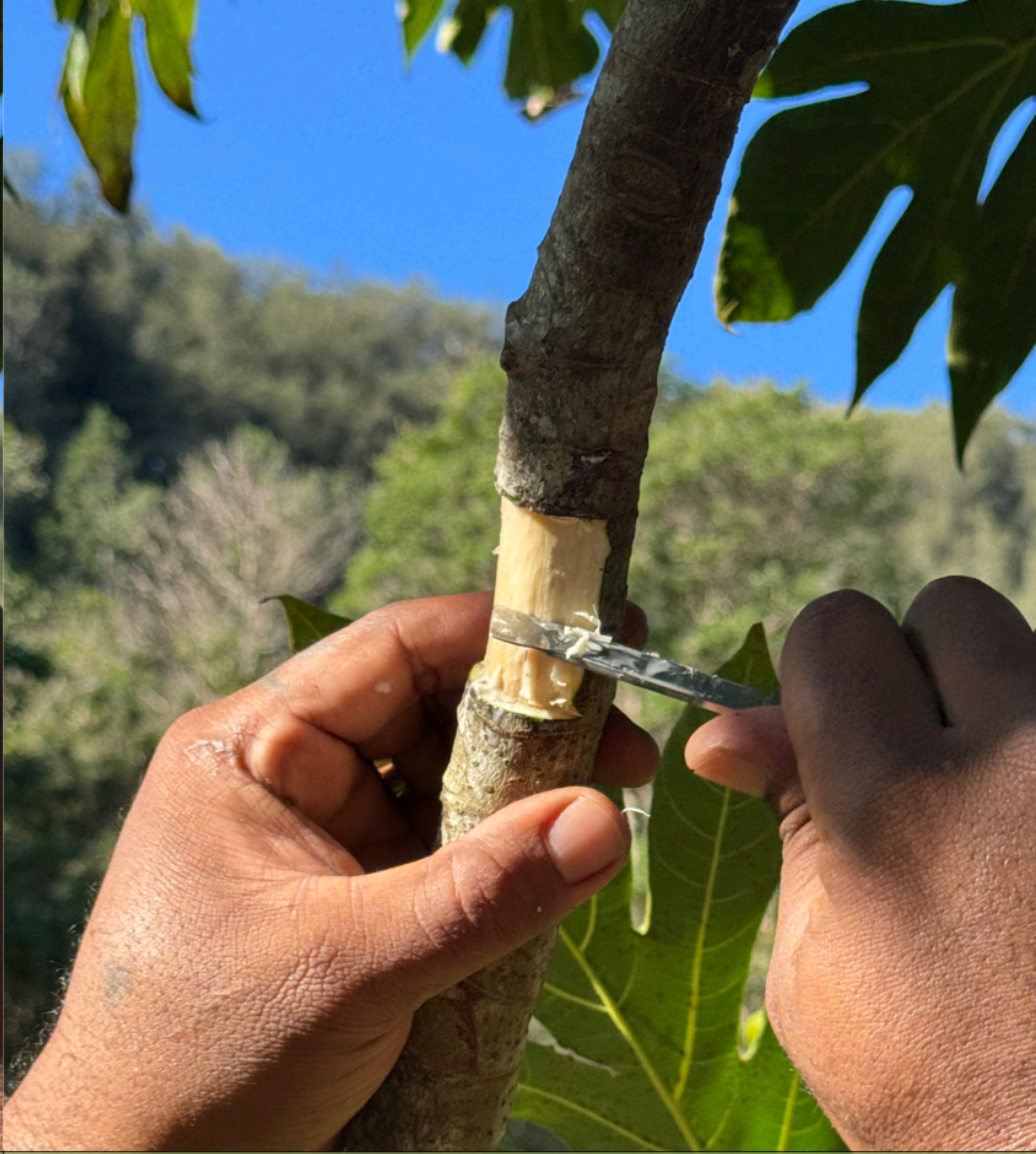

| Step | Procedure | Key considerations |
|---|---|---|
| 2 | Remove 2–3 cm of bark and cambium from the circumference of targeted internodal section of branch, exposing xylem. | Ensure bark is removed from the entire circumference of the targeted, internodal section. |

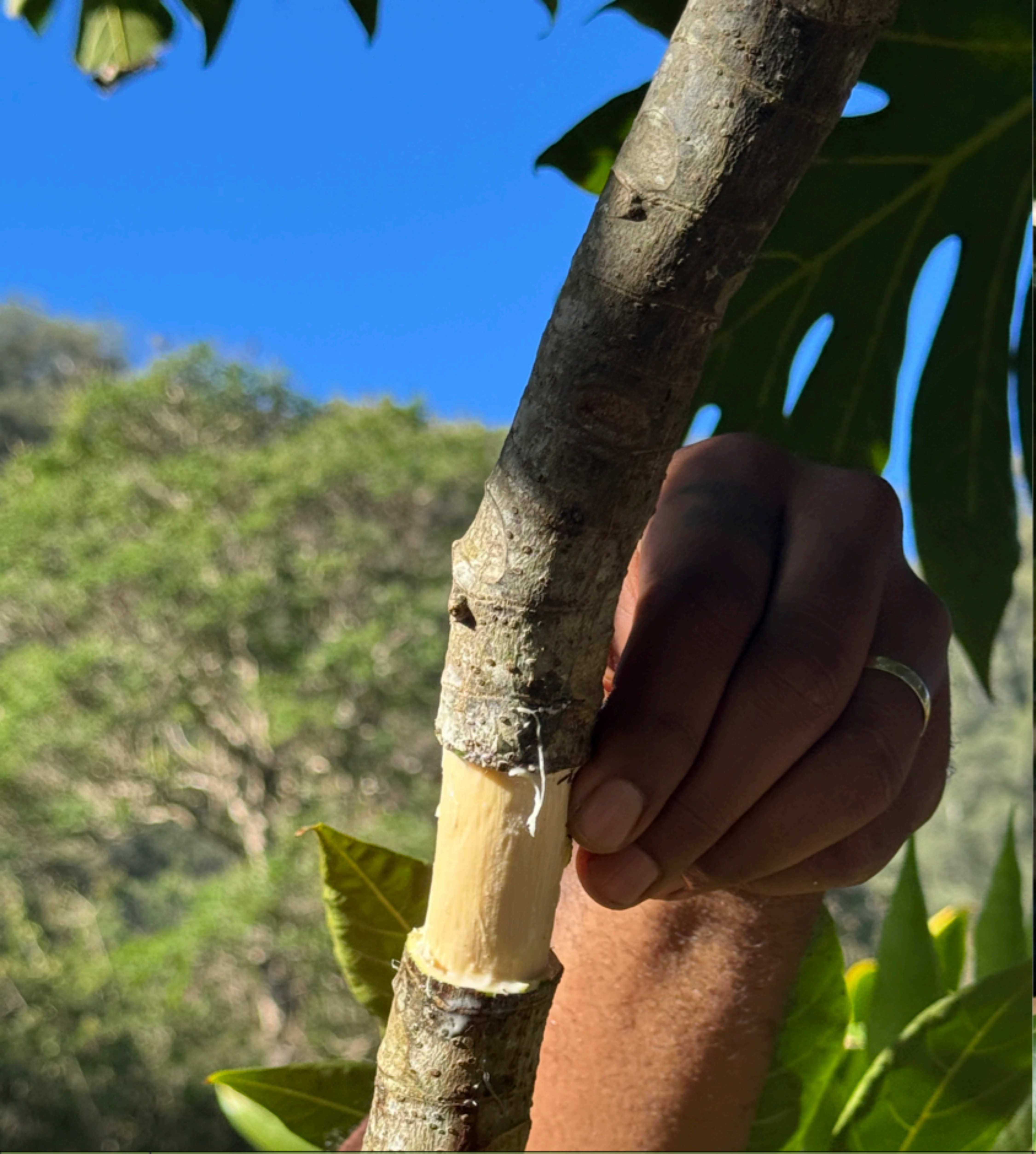

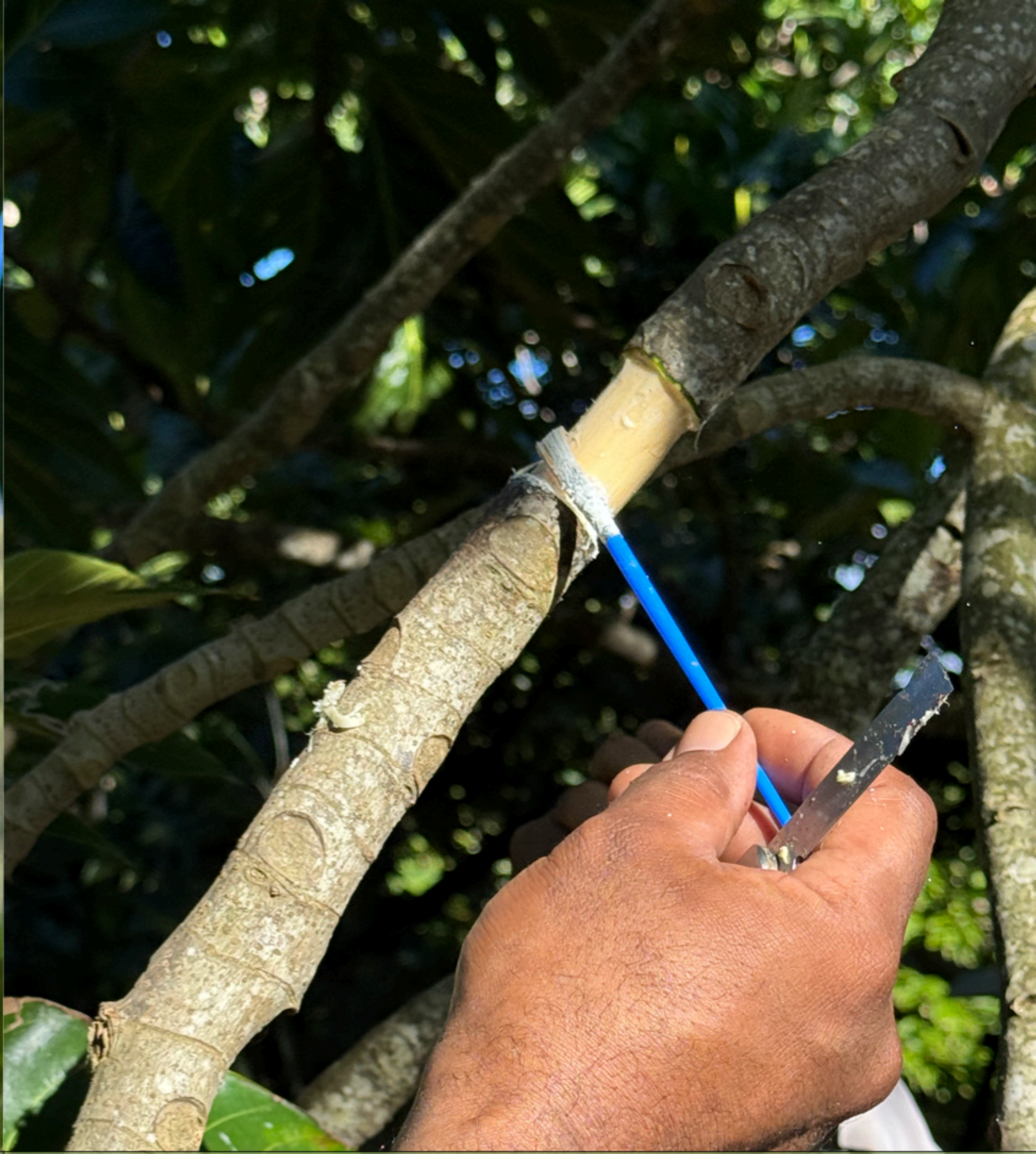

| Step | Procedure | Key considerations |
| --- | --- | --- |
| 3 | With blade at 90° angle, scrape exposed xylem lightly ensuring no cambial tissue remains. Apply rooting hormone to girdled node. | Complete cambial tissue removal is important for preventing callus bridging (wound healing, bark regrowth). Hormone application optional. |

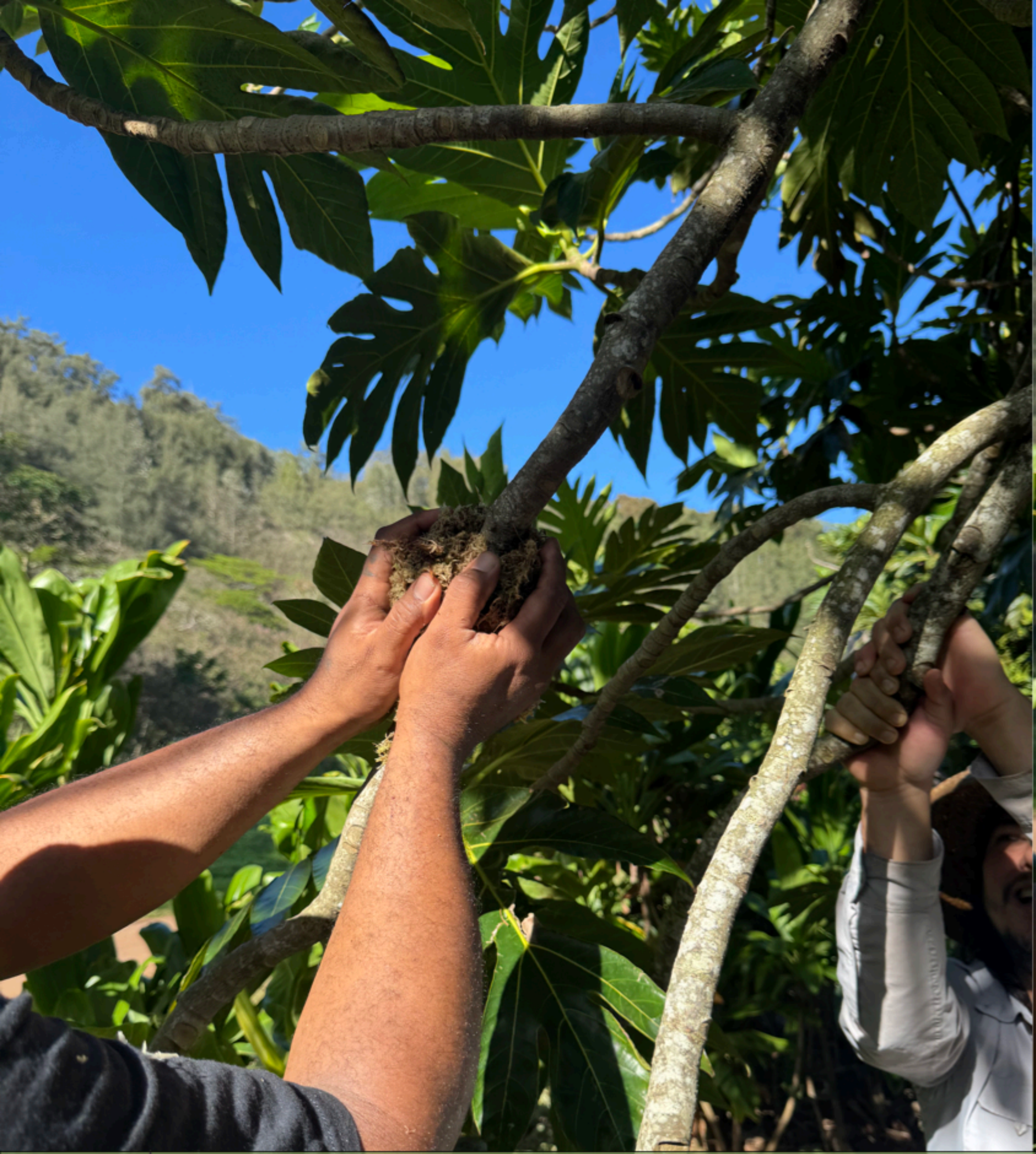

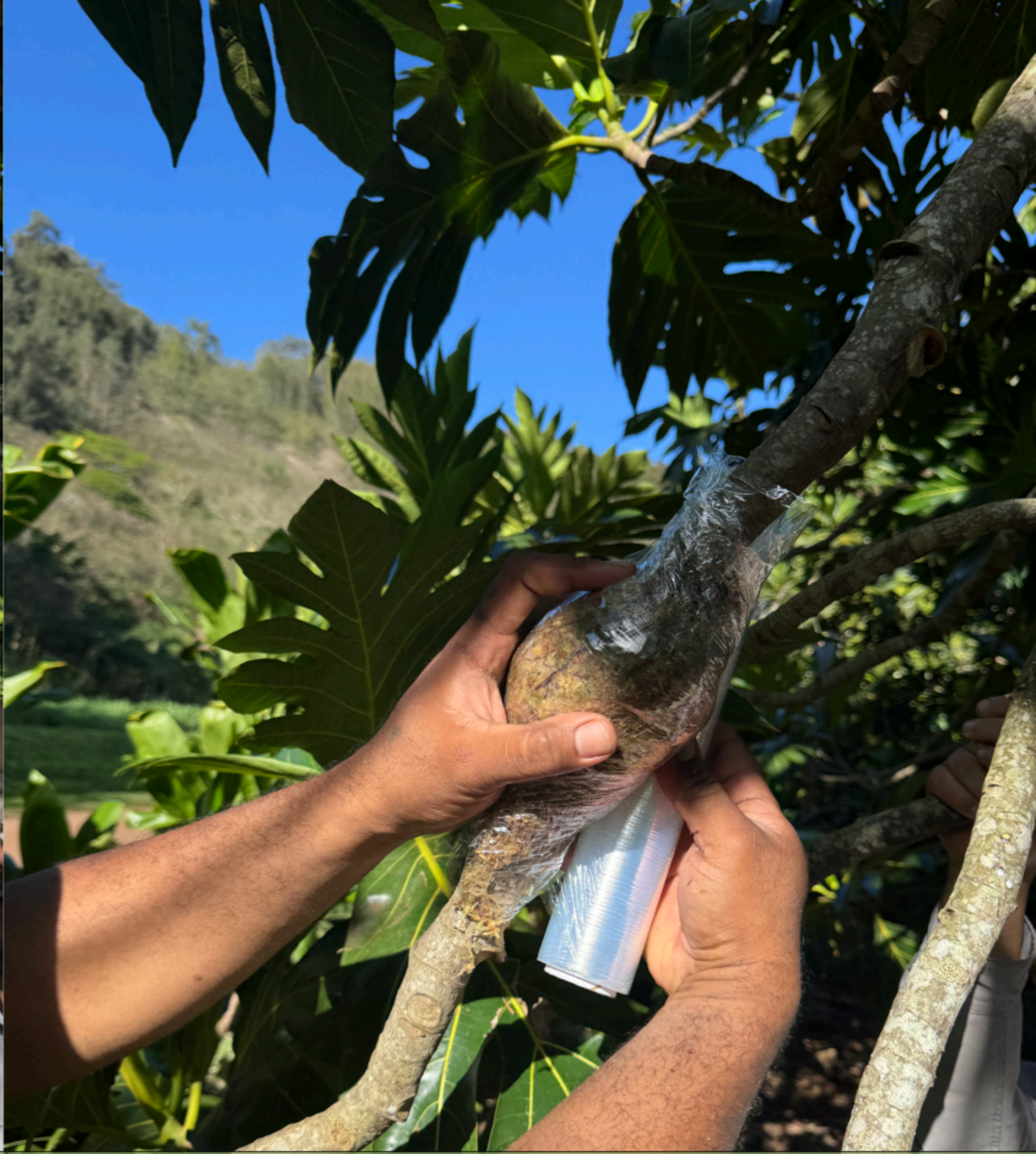

| Step | Procedure | Key considerations |
| --- | --- | --- |
| 4 | Soak and apply moist rooting media evenly around girdled area and enclose with plastic wrap. | Media e.g. coco coir or sphagnum moss. Squeeze excess moisture from media which should be damp, not saturated to prevent rot. |

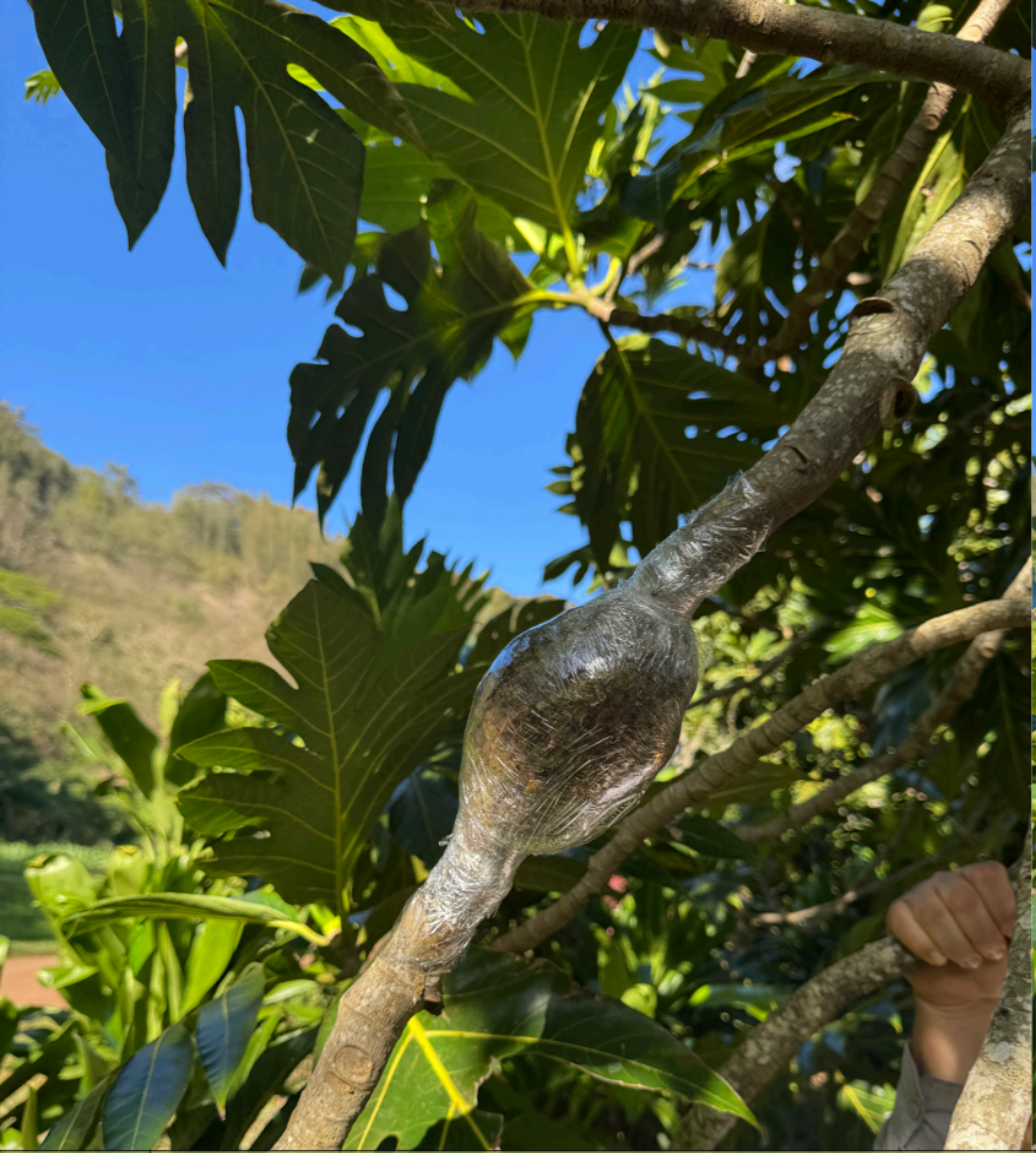

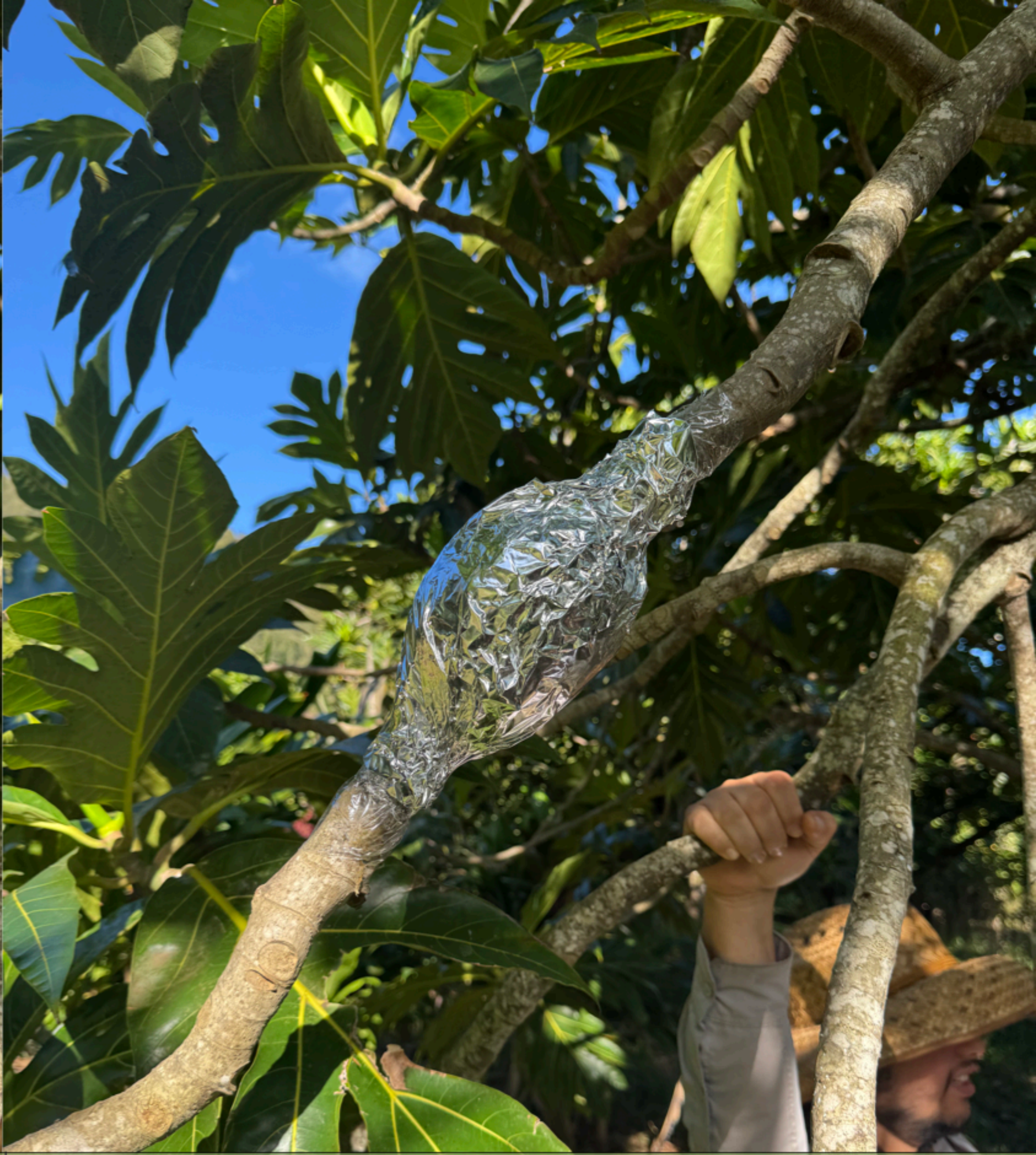

| Step | Procedure | Key considerations |
|---|---|---|
| 5 | Secure plastic tightly above and below girdle. Cover wrapped air layer with opaque material (i.e., aluminum foil). Monitor bi-weekly, harvest once root development stabilizes. | The covering excludes light, promoting root initiation. Attach label (date, name, variety ID, tree ID). Time to root development varies, ideally percent root coverage at harvest >50%. |
